# A SLiM-dependent conformational change in baculovirus IE1 controls its focus formation ability

**DOI:** 10.1101/2023.03.06.531455

**Authors:** Toshihiro Nagamine, Yasushi Sako

## Abstract

The baculovirus IE1 gene encodes a multifunctional protein that is essential for both DNA replication and RNA transcription of the virus. Prior to viral DNA replication, IE1 promotes early gene transcription when localized in *hr*-dependent foci. During viral DNA replication, the IE1 foci expand and fuse to generate the virogenic stroma (VS) with IE1 forming the VS reticulum. To explore the IE1 structural features essential for this coordinated localization, we constructed various IE1 mutants based on three putative domains (N, I, and C). We determined that a BDI motif located in the intrinsic disorder region (IDR) between the N and I domains acts as a nuclear localization signal, whereas BDII and HLH in the C domain are required for VS localization in infected cells or for chromosomal association in uninfected mitotic cells. Deletion of the SLiM (<u>s</u>hort <u>li</u>near <u>m</u>otif) located in the I domain restrains both nuclear- and VS localization. Intra-molecular fluorescence resonance energy transfer (FRET) probes of IE1 mutants revealed a conformational change of the I-C two-domain fragment during infection, which was inhibited by aphidicolin, suggesting that IE1 undergoes a stage-dependent conformational change. Further, homo-dimerization of the I domain and stage-dependent conformational changes require an intact SLiM. Mutational analysis of SLiM revealed that VS localization and chromosomal association were retained following S291A and S291E substitutions, but *hr*-dependent focus formation differed between the two mutations. These results suggest that coordinated IE1 localization is controlled by SLiM-dependent conformational changes and that are dependent on the SLiM phosphorylation state.

**Importance:** SLiMs (<u>s</u>hort <u>li</u>near <u>m</u>otifs) are compact non-globular protein interaction interfaces that mediate various cellular functions and as such are often mimicked by viruses to rewire cellular pathways. Here, we found that an unusual type of viral SLiM acts as a conformational switch rather than a cellular mimic. Prior to viral DNA replication, the baculovirus IE1 protein promotes early gene transcription within its focus. During viral DNA replication, the IE1 foci expand and fuse to form the virogenic stroma where late virus replication events occur. Our results indicate that IE1 undergoes a conformational change that is dependent on the infection stage and SLiM phosphorylation. This work provides new insights into the role SLiMs play in heterotypic and homotypic protein-protein interactions.

## Introduction

Baculoviruses are large insect-pathogenic DNA viruses that are widely employed as expression vectors for recombinant protein production (1). In addition to their practical importance, baculoviruses are interesting targets for biological research in terms of their unique replication mechanism and long co-evolutional history with host insects (1, 2). Baculovirus genes are expressed in a sequential and coordinated manner during infection. Once the baculovirus genome enters the nucleus, immediate early genes including *ie1* are expressed without the support of other viral genes (1). Subsequently, prior to viral DNA replication, IE1 promotes delayed early gene transcription within its focus distribution (3, 4). These early transcription products then serve in viral DNA replication and late gene transcription. During DNA replication, the IE1 foci expand to form the virogenic stroma (VS), a substructure of which is the VS reticulum in which IE1 associates with viral DNA, while other viral proteins are concomitantly localized to DNA-sparse regions (5). The IE1 intracellular localization thus varies depending on the stage of infection; however, the IE1 structural features involved in this coordinated localization have yet to be fully elucidated.

Previous reports have suggested that a number of specific motifs (short sequences) are involved in IE1 nuclear import and DNA binding. BDI [Basic domain I; residues 152 to 161 in the Autographa californica multiple nucleopolyhedrovirus; (AcMNPV)] was originally identified as a highly conserved domain required for IE1 interactions with the *hr* enhancer element and subsequent formation of an *hr*-dependent focus (6, 7). A recent report using a nuclear localization signal (NLS) prediction algorithm identified a putative bipartite NLS in BDI (8, 9), which we propose to subdivide into BDIa and BDIb with the original BDI re-defined BDIb as shown in Figure 1A. Additional stretches of basic amino acids encompassing the C termini of BDII (residues 521 to 538 in AcMNPV) and HLH (Helix-loop-helix dimerization motif; residues 543 to 568 in AcMNPV) play roles in DNA binding and nuclear import (10, 11). Although the existence of these motifs has been known for a long time, the relationship between these motifs and the coordinated localization of IE1 is not well understood.

**Fig. 1.**
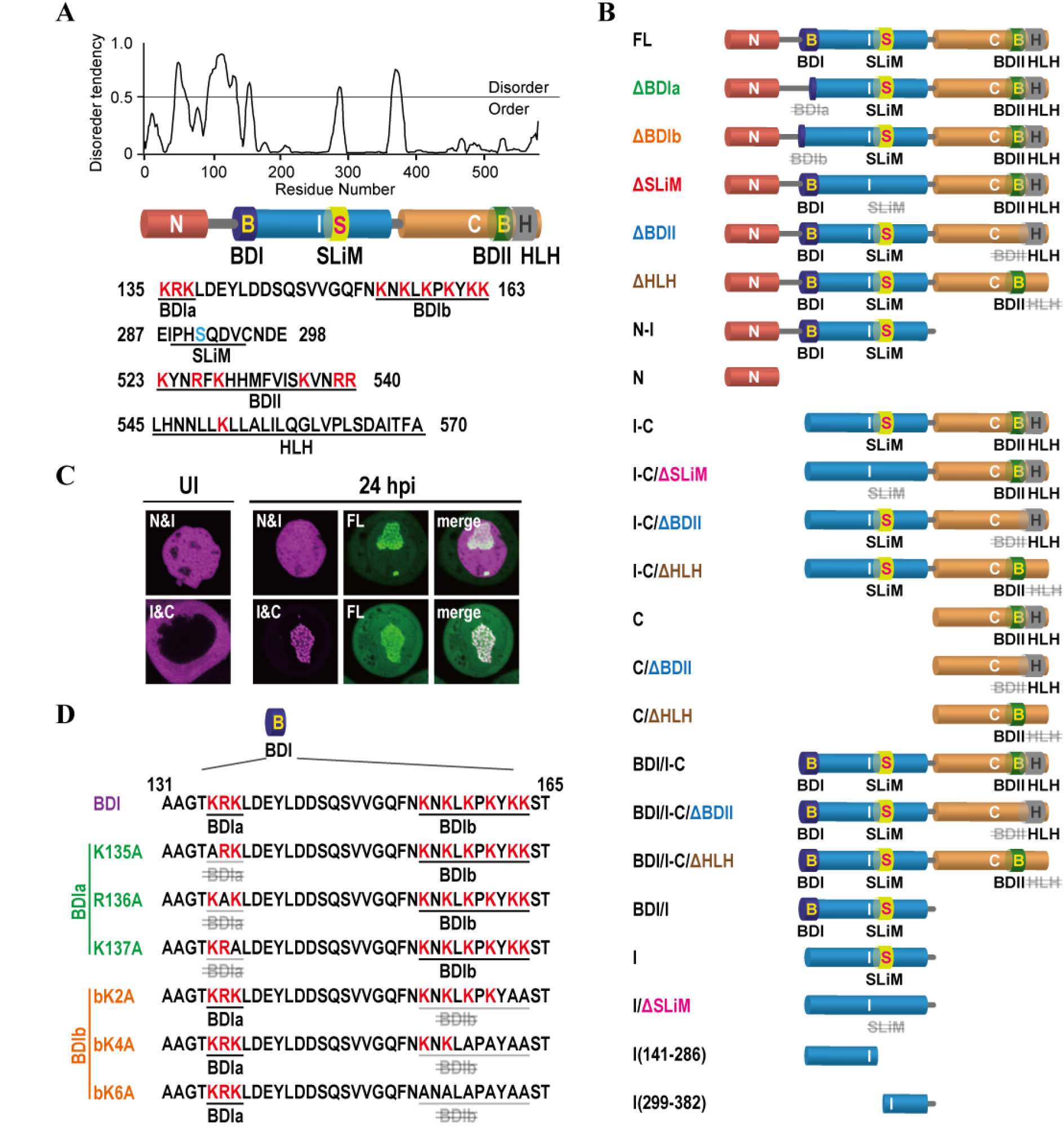
Prediction of a three domain architecture in IE1 and schematic representation of the experimental fragments assayed. (A) PONDR score for the IE1 sequence and the IE1 motif sequences. Red letters indicate basic amino acid residues. The blue letter indicates the serine residue (S291) in the SLiM. (B) Schematic representations of the IE1 fragments assayed. (C) Intracellular localization of the N-I and I-C fragments in uninfected (UI) and infected cells (24 hpi). (D) BDI and associated mutant sequences. Red letters indicate basic amino acid residues.

SLiMs (short linear motifs) are compact non-globular protein interaction interfaces that mediate a variety of cellular functions, including targeting proteins to specific subcellular compartments, providing specificity for post-translational modifications, and serving as binding sites for components of multiple regulatory and signaling pathways (12). For viral replication to proceed various cellular functions need to be modulated; however, constraints on virus genome sizes limit encoded options. To overcome this limitation, viruses hijack cell regulation by mimicking cellular SLiMs (13). In contrast to this cellular mimic role, we have identified an atypical type of viral SLiM that functions as a conformational switch. Here, we show that the baculovirus IE1 SLiM is required for domain interactions and conformational changes in IE1. While two SLiM mutations (S291A and S291E) retained VS localization and chromosomal association, their effects on *hr*-dependent focus formation differed. These results suggest that the cooperative function among IE1 motifs underlying IE1 localization is controlled by SLiM-dependent conformational changes that are potentially switched by its phosphorylation states.

## Results

### IE1 is expected to consist of three domains based on a disorder tendency score

To investigate the IE1 structural features required for its coordinated localization, we generated GFP-tag or Halo-tag (labeled with TMR: Tetra-methyl-rhodamine) constructs of BmNPV (Bombyx mori nucleopolyhedrovirus) IE1 fragments and examined their subcellular localization. As the baculovirus IE1 molecular structure has not been determined, we predicted a putative tertiary domain architecture consisting of N (1-100), I (141-382), and C (369-581) domains based on a disorder tendency score (PONDR) (Fig. 1A). We initially examined the subcellular localization of two-domain fragments, N-I and I-C (Fig. 1B and C). In uninfected cells, N-I exhibited a nuclear localization similar to wild-type IE1 (FL), and I-C localized to the cytoplasm (Fig. 1C and see Fig. 2A). In contrast, in infected cells at 24 hpi, I-C localized to the VS similar to FL, whereas N-I did not exhibit VS localization (Fig. 1C). We hypothesized that the localization patterns were motif dependent and investigated each of the motifs.

**Fig. 2.**
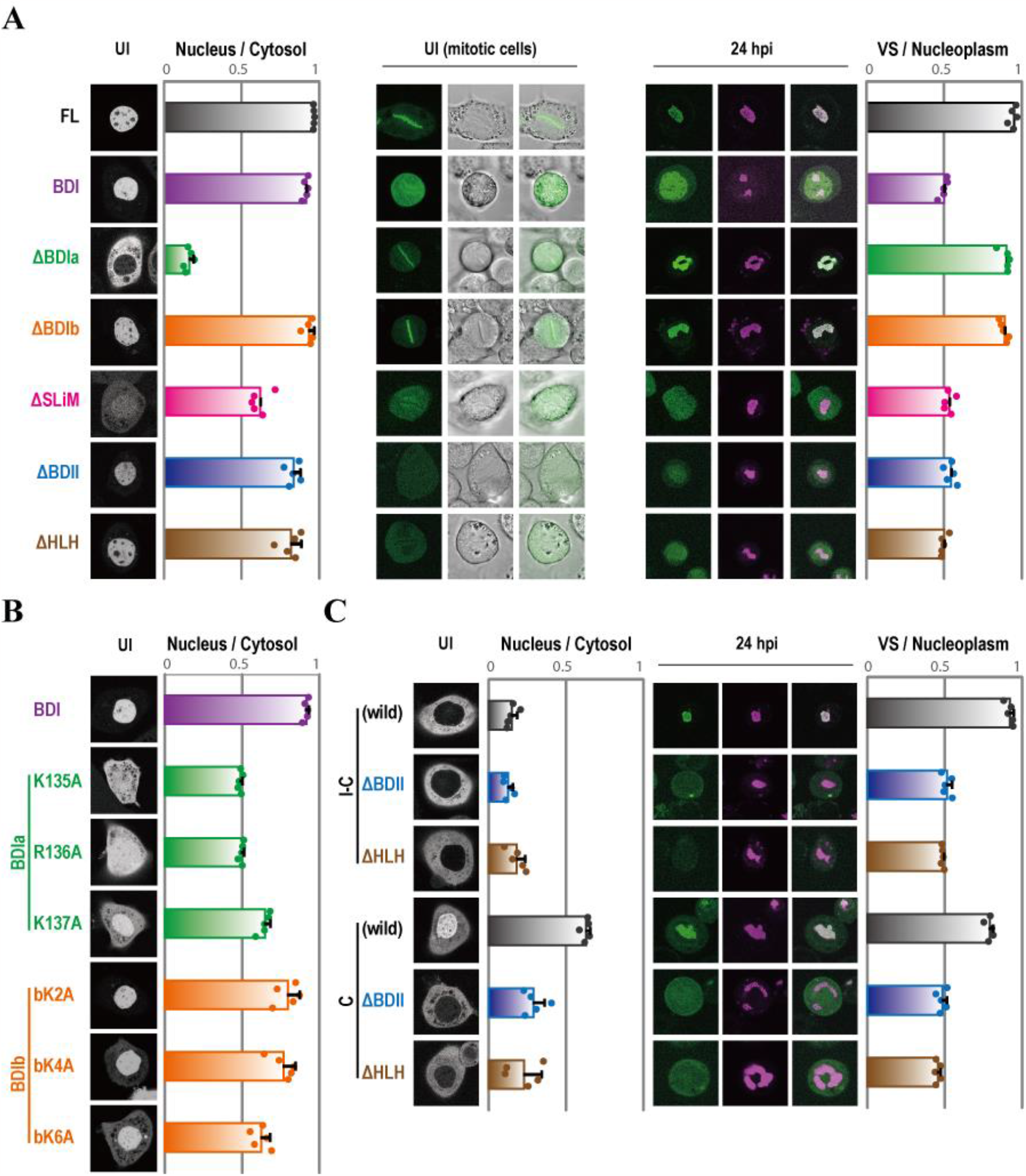
Intracellular localization of IE1 mutants in uninfected and infected cells. (A) Distribution ratio of nucleus / cytosol localization of the FL mutants in uninfected cells (UI) and VS / nucleoplasm localization in infected cells (24 hpi). (B) Nucleus / cytosol localization ratio of the BDI mutants in uninfected cells. (C) Distribution ratio of nucleus / cytosol localization of the I-C and C mutants in uninfected cells (UI) and VS / nucleoplasm localization in infected cells (24 hpi). The indices are represented as the mean ± SD (error bars) and points on the graph. n = 5.

### BDI functions as a nuclear localization signal

The original BDI (Basic domain I; residues 152 to 161 in AcMNPV) is a highly conserved domain required for IE1 binding to the *hr* enhancer element as well as *hr*-dependent transactivation and focus formation (6, 7). BDI was recently predicted (8, 9) to comprise a bipartite NLS (Fig. 1A). We thus subdivided the motif into BDIa (135-137) and BDIb (154-163) and renamed the original BDI as BDIb (Fig. 1A). Since I-C has BDIb but lacks BDIa (Fig. 1B), the roles of the motifs were analyzed separately. While a full-length mutant lacking BDIb (ΔBDIb) localized to the nucleus, the reciprocal mutant lacking BDIa (ΔBDIa) did not exhibit nuclear localization (Fig. 1B and 2A). This suggests that BDIa, rather than BDIb, is required for the nuclear localization of IE1. If so, BDIa alone should be sufficient as a functional NLS. A prediction algorithm (8) indicated that the bipartite NLS in BDI (containing both BDIa and BDIb) includes three basic amino acids in BDIa (Fig. 1D). Consistent with this prediction, replacement of these amino acids impacted nuclear localization (Fig. 2B). In contrast, replacement of two or four of the six basic amino acids in BDIb (Fig. 1D) had little effect on nuclear localization, whereas the absence of all six resulted in an ambiguous nuclear localization pattern (Fig 2B). This indicates that BDIa alone is insufficient as an independent NLS. Since the full-length mutant ΔBDIb exhibited an explicit nuclear localization pattern, the DNA binding activity of BDII and HLH may facilitate nuclear localization of ΔBDIb (see below).

### BDII and HLH are responsible for DNA binding

BDII (residues 521 to 538 in AcMNPV) and HLH (residues 534 to 568 in AcMNPV) have been shown to play a role in DNA binding and nuclear import (10, 11). To confirm this, we constructed full-length mutants lacking BDII or HLH (ΔBDII and ΔHLH) and examined their subcellular localization. In uninfected (inter-phase) cells, both of the mutants localized to the nucleus, probably due to BDI function; however, unlike FL (a wild-type IE1), neither localized to the VS in infected cells at 24 hpi nor were they associated with cellular chromosomes in uninfected mitotic cells (Fig. 1B and 2A). In contrast, ΔBDIa and ΔBDIb localized to the VS and cellular chromosomes. These results suggest that BDII and HLH are required for DNA binding and their DNA binding activity may contribute to VS localization. To confirm that BDII and HLH function in VS localization, the subcellular distribution of I-C- and C-fragments lacking BDII or HLH was also examined. Both domains were required for the VS localization of the I-C- and C-fragments as well as FL (Fig. 2 A and C). Notably, whereas the C-fragment retained BDII- and HLH-dependent nuclear localization in uninfected cells, this was absent in the I-C-fragment (Fig. 2C). This suggests that the I domain masked nuclear retention of the C-fragment.

### I-C undergoes a BDII- and HLH-dependent conformational change during infection

To address the I domain function, we next examined conformational changes between the FL and various fragments in infected and uninfected cells using intra-molecular fluorescence resonance energy transfer (FRET) probes. The FRET probes were constructed by fusing GFP and Halo-tag (TMR) at the N- and C-termini respectively of the target sequences. The FRET index was determined by measuring the TMR / (GFP + TMR) fluorescence intensity ratio following stimulation with a 488 nm-laser (for GFP). For the FL-FRET probe, there was no significant difference in the FRET indices between infected and uninfected cells due to relatively diverse values among individual infected cells (Fig. 3A). This diversity might be attributable to different sub-phases of the late infection stage as the cells were measured at 24 hpi (see Discussion). In contrast, the FRET indices of the N-I and I-C fragments were significantly different between infected and uninfected cells. Since the FRET indices of the single-domain N fragment likewise differed between infected and uninfected cells, the effects seen with N-I fragment could be N fragment dependent (Fig. 3A). Although the I-C fragment had different FRET indices, the I and C single-domain values were stable regardless of infection status (Fig. 3A). This suggests that the spatial arrangement of the I and C fragments changes during infection. The FRET indices of BDII- or HLH-deleted I-C fragments were similar in infected and uninfected cells, suggesting that BDII and HLH contribute to the conformational change (Fig. 3B). In addition, an I-C fragment bearing BDI (BDI/I&C:131aa-581aa) had similar values as the cytoplasmic I-C (141aa-581aa) in infected and uninfected cells, suggesting that the intracellular location of I-C does not affect its spatial arrangement. (Fig. 3B). Aphidicolin, a DNA synthesis inhibitor, blocks viral DNA replication and VS formation. In cells treated with this reagent, the FRET indices of BDI-I-C in IE foci (mean 0.54) were similar to those in uninfected cells (mean 0.53) at 24 hpi, suggesting that the conformational change occurs in the late stages of infection (Fig. 3C).

**Fig. 3.**
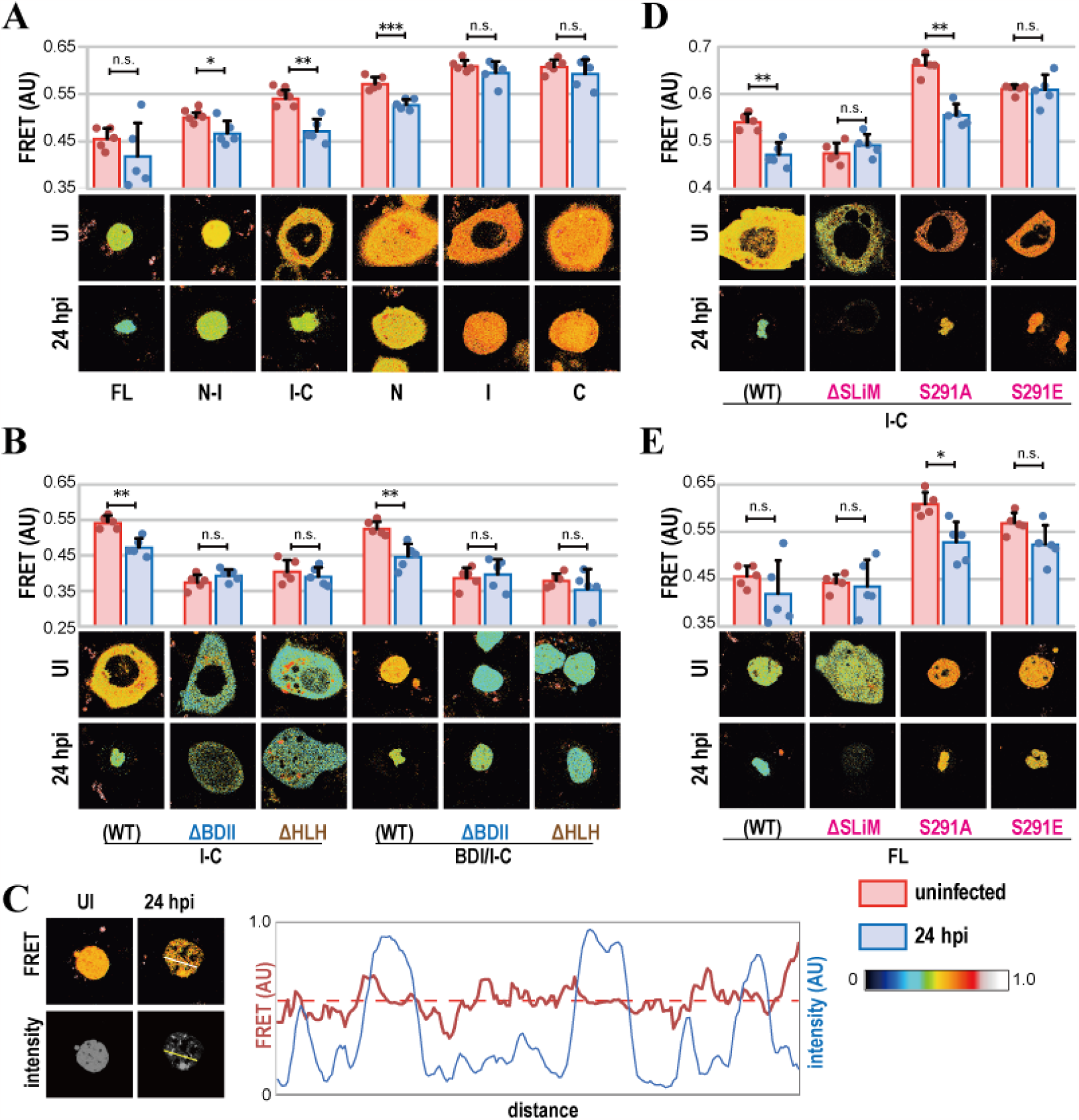
Conformational comparison of IE1 mutants in uninfected (UI) and infected cells (24 hpi) using FRET probes. FRET differences were determined by measuring the TMR / (GFP + TMR) fluorescence intensity ratio of the IE1 fragments (A), the BDII- and HLH-deletion mutants of I-C and BDI/I-C (B), I-C in aphidicolin-treated cells (C), I-C SLiM mutants (D) and FL (E) when probes were irradiated with a 488 nm-laser (for GFP). The line graph shows the FRET index and fluorescence intensity on the yellow line in aphidicolin-treated and infected cells (24 hpi) in (C). For (A, B, D, and E) FRET indices are represented as the mean ± SD (error bars) and points on the graph. n = 5. *p* values were calculated using paired t test: \**p* < 0.05, \*\**p* < 0.01, \*\*\**p* < 0.001, n.s., not significant.

### A SLiM candidate is required for the I-C conformational change

SLiMs (short linear motifs) are compact non-globular protein interaction interfaces that mediate a variety of cellular functions, including targeting proteins to specific subcellular compartments such as NLS and providing specificity for post-translational modification (12). We identified a SLiM candidate in the I fragment (Fig. 1A). To investigate the function of this sequence, we initially constructed a full-length mutant in which the motif was deleted and observed its subcellular localization (Fig. 2A). In uninfected (inter-phase) cells, the mutant exhibited a vague nuclear localization pattern, suggesting that the SLiM candidate affects BDI function. Failure to localize to the VS in infected cells and absence of chromosome binding in uninfected mitotic cells suggest that the motif plays a role in DNA binding. To determine if it is involved in the I-C conformational change, we next constructed FRET probes of I-C-fragment mutants and full-length mutants with deletions or single point mutations in the SLiM sequence. Deletion of SLiM in I-C disturbed the conformational change of the fragment and decreased its expression, possibly due to instability (Fig. 3 D and E). SLiMs are frequently targets for post-translational modifications. Consistent with this, the putative SLiM we identified has a potential phosphorylation site (S291) in the middle of the sequence (Fig. 1A). We thus mutated S291 to alanine or glutamic acid, a mimic of phosphorylated serine. The FRET indices of I-C/S291A were significantly different between infected and uninfected cells and were higher than those of the unmutated I-C, suggesting that I-C/S291A undergoes a conformational change similar to that of unmutated I-C during infection (Fig. 3D). In contrast, no significant differences were observed between infected and uninfected cells with I-C/S291E, suggesting that it remains stable during infection (Fig. 3D). These results imply that the I-C conformational change requires an unphosphorylated S291. The full-length mutants behaved similarly to the I-C-fragment mutants except that there were no significant differences for FL (a wild-type) between infected and uninfected cells (Fig. 3E).

### The SLiM plays a role in homo-dimerization of the I domain

Because SLiMs are protein interaction interfaces, we sought to determine the interaction partner(s) of the identified SLiM candidate. When analyzing the IE1 dimerization mechanism, we noticed possible dimerization of the I-domain by itself (see below). To determine if the SLiM region is involved, we co-expressed experiments I fragments containing BDI (BDI/I) and various I mutants lacking BDI. While the nonmutated I fragment (without BDI) localized to the cytoplasm in single-transfected cells, its nuclear distribution increased in BDI/I co-transfected cells (Fig. 4A). This suggests that dimerization is associated with the I domain. Further, deletion of SLiM in the I fragment resulted in cytoplasmic localization even in BDI/I co-transfected cells, suggesting that the dimerization critical region had been lost. Mutation of the putative S291 phosphorylation site (S291A or S291E) in the SLiM region of the I fragment did not disrupt dimerization. We speculated that the SLiM interaction partner would consist of an ordered region, two of which are present in amino acids 141-286 and 299-382 of I (Fig. 1B). Although the I(299-382) fragment did not interact with BDI/I, the I(141-286) fragment did, albeit on a more limited basis than that observed with the full BDI/I (Fig. 4A). These results suggest that the putative SLiM is functional and that homo-dimerization of the I fragment is directly mediated by two interaction points between SLiM and I(141-286). Considering the two interactions, the I-domain dimer conformation is likely antiparallel (see Fig. 1A).

**Fig. 4.**
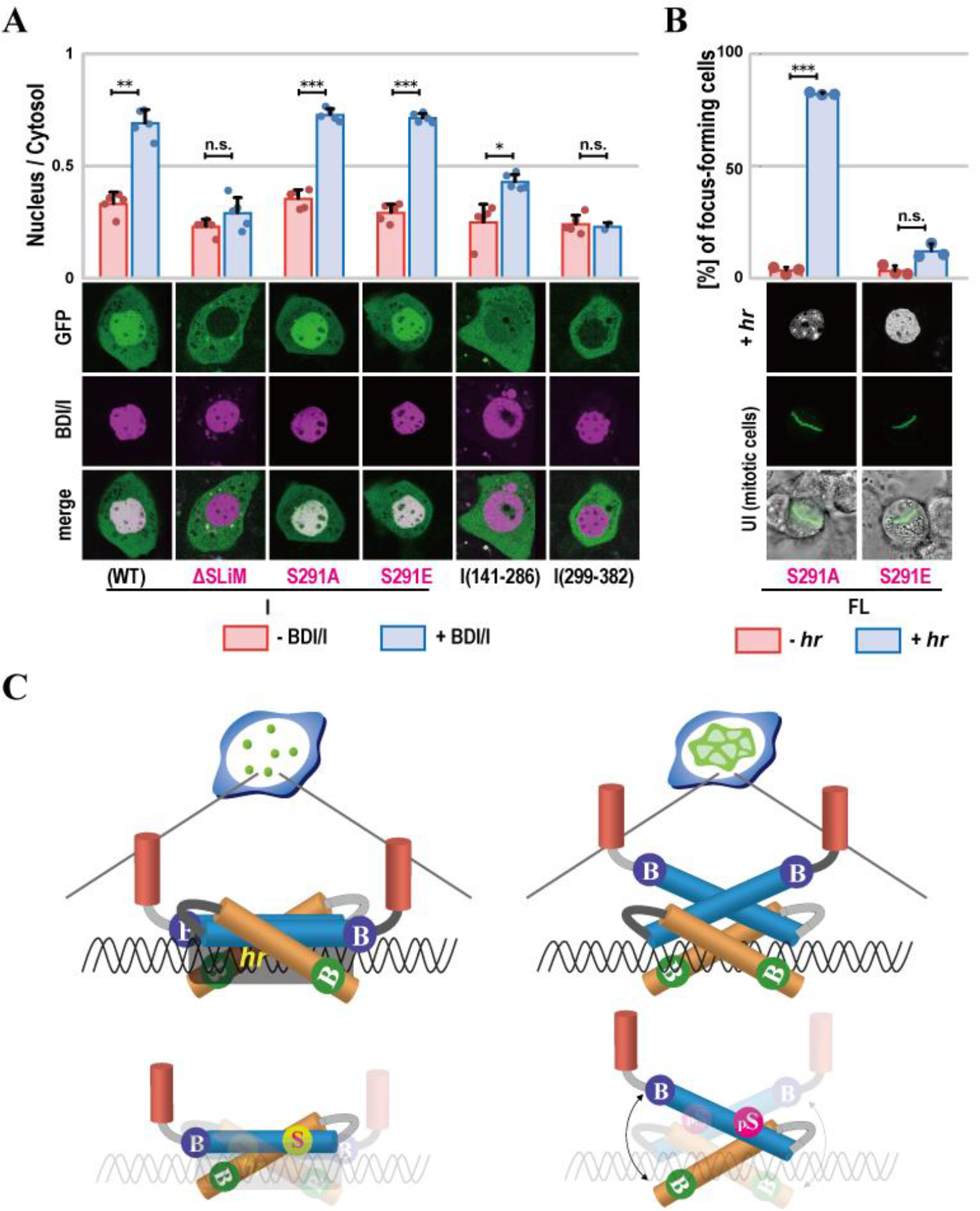
The IE1 SLiM plays a role in homo-dimerization of the I domain and IE1 *hr*-dependent focus formation. (A) Distribution ratio of nucleus / cytosol localization of I in single-transfected (-BDI/I) and BDI/I co-transfected cells (+ BDI/I). (B) Percentages of focus-forming cells transfected with FL SLiM mutants in the absence (- *hr*) and presence of *hr* (+ *hr*). (C) Schematic drawing of the different IE1 conformations. While in the early stages of infection (left) IE1 binds *hr* via a compact conformation with S291unphosphorylated, in the late stages of infection (right) IE1 binds to sequence-nonspecific DNA via an extended conformation with S291phosphorylated. For (A and B) the values are represented as the mean ± SD (error bars) and points on the graph. n = 5. *p* values were calculated using paired t test: \**p* < 0.05, \*\**p* < 0.01, \*\*\**p* < 0.001, n.s., not significant.

### A phosphorylated mimic of SLiM (S291E) perturbs *hr*-dependent focus formation

SLiM involvement in the I-C conformational change, possibly via its interaction with I(141-286), seems to be regulated by its phosphorylation state during infection. In response to the structural change, the localization of IE1 shifts from a focus distribution to association with the VS reticulum. To explore this relationship between conformation and localization, we examined two SLiM mutants, FL/S291A and S291E, for effects on *hr*-dependent focus formation. Both mutants localized to uninfected mitotic chromosomes, suggesting that they remain capable of binding DNA (Fig. 4B). FL/S291A also retained *hr*-dependent focus formation activity (Fig. 4B). In contrast, focus formation was lost with the phosphorylated SLiM mimic, FL/S291E (Fig. 4B). These results suggest that IE1 with an unphosphorylated SLiM can form *hr*-dependent foci and that SLiM phosphorylation leads to IE1 association with the VS reticulum depending on sequence-nonspecific DNA binding rather than *hr*-specific binding.

## Discussion

In the early stages of infection, it seems that the localization of IE1 to *hr*s in parental (entrant) virus DNA depends on the binding of BDIb with *hr*s because mutations in BDIb perturbs IE1 binding to *hr in vitro* and focus formation in uninfected cells (6, 7). It is thus likely that the *hr*-specific DNA binding of IE1 serves to discriminate viral DNA from cellular DNA. The IE1 focus containing the parental virus DNA can then recruit RNA transcription factors and DNA replication factors needed for a shift to late stage infection, when the IE1 foci expand and form the VS reticulum. This VS structure consists of two sub-structures, a DNA-rich region and a DNA-sparse region, that may contribute to the DNA packaging process, in which progeny virus DNA is inserted into capsids at the interface of the two regions. Thus VS reticulum formation may require sequence-nonspecific DNA binding rather than *hr*-specific binding by IE1, which is one possible reason for the IE1 conformational change illustrated in Figure 4C.

FL/S291Aretained both the I-C conformational change and *hr*-dependent focus formation. However, changing the mutation to S291E, which mimics a phosphorylated SLiM IE1, resulted in a loss of both properties despite maintaining sequence-nonspecific DNA binding (Fig. 4B). These findings suggest that the phosphorylation state of SLiM in IE1 affects its conformation and DNA-binding properties. In the early stages of infection, IE1 with an unphosphorylated SLiM adopts a compact conformation (high I-C FRET state), in which BDIb, BDII, and HLH may contribute cooperatively to *hr* binding because loss of these motifs reduces the FRET indices of I-C (Fig. 3 B and D). It is possible that the compact conformation of IE1 with unphosphorylated SLiM suppresses the sequence-nonspecific DNA binding of BDII and HLH. In the late stages of infection, on the other hand, it is likely IE1 SLiM phosphorylation (by a viral or virus-activated cell kinase) promotes a conformation change in IE1 from the compact form to an extended form (low I-C FRET state). This conformational change would alter the DNA binding property of IE1 from *hr*-specific to sequence-nonspecific DNA binding, possibly to form the VS reticulum. However, I-C/S291A undergoes the presumed conformational change despite the absence of phosphorylation. Although the reasons for this are unclear, one possibility is that I-C/S291A may heterodimerize with intact IE1 in virus-infected cells. If so, the SLiM in the intact IE1would be phosphorylated and the phosphorylated SLiM could lead to the extended conformation even though SLiM in I-C/S291A is unphosphorylated. This ability to change the structure with a single phosphorylation event may allow for quick responses. It is also unclear mechanistically how the SLiM phosphorylation state drives I-C conformational changes. Whereas the FRET indices of I-C (WT) and I-C/S291A were affected by viral infection, their respective values differed. This suggests that S291 and A291 differently influence the I-C conformation in infected and uninfected cells. Since the IE1 SLiM plays a role in anti-parallel homo-dimerization of the I domain, the strength of the I-domain homo interaction may be important for the I-C conformation.

The N domain and the downstream intrinsic disordered region (IDR; 97-140) have long been known as the acidic activation domain (AAD) that is required for both transcriptional activation and DNA replication and which is phosphorylated during infection (14, 15). Both N-terminal domains are structurally more flexible than I and C (Fig. 1A) and are unlikely to play a role in DNA binding. Rather, these domains are thought to be involved in heterotypic protein interactions. It is possible that the differences in the values of the N-domain FRET indices between uninfected and infected cells and those from diverse FL constructs in individual infected cells could be due to variations in the phosphorylation state and protein-protein interactions of the N-terminal N and IDR domains. Further analysis of the interaction between IE1 and other proteins is needed to better elucidate the IE1structure-function relationship in more detail.

In conclusion, we have proposed a three-domain structure (N, I, and C domains) for IE1 based on a disorder tendency score (PONDR) and shown a change in the spatial arrangement between the I and C domains during infection. This conformational change in I-C may be regulated by the phosphorylation state of the SLiM in the I domain, which could switch IE1 DNA binding from *hr*-specific to sequence-nonspecific. While most cellular and viral SLiMs contribute to heterotypic protein-protein interactions (12, 13), the baculovirus SLiM characterized here appears to be involved in anti-parallel homo-dimerization of the I domain, the strength of which may affect the I-C conformational state.

## Materials and methods

### Cells and viruses

BmN cells were maintained in TC100 medium (AppliChem) supplemented with 10% FBS (4). The BmNPV wild-type isolate T3 (16) was propagated in BmN cells.

### Plasmid construction

The plasmids expressing GFP-tagged- and Halo-tagged IE1 fragments were constructed from pHE-C, a pEGFP-C1 (Clontech)-derived plasmid (5), and pHH-C (17), a pHE-C-derived plasmid, respectively. The plasmids expressing the FRET probes were derived from phsp-GFP-Halo, which was constructed by inserting the HaloTag open reading frame of pHalo7-C1 (17) into the BamHI site of pHE-C. DNA fragments encoding the proteins of interest were PCR amplified (KOD One PCR Master Mix;Toyobo) with oligonucleotides and templates listed in Table S1 and inserted into the SalI site of pHE-C, pHH-C, or phsp-GFP-Halo by a seamless cloning method with SLiCE (18) (Table S1). For construction of deletion mutants, inverse PCR amplification was conducted with oligonucleotides and templates listed in Table S1 and the resulting PCR products were treated with T4 Polynucleotide Kinase (Takara) and ligated with a DNA ligation kit (Takara). The constructs expressing GFP-BDI-bK6A and single amino acid substitutions in the SLiM sequence were generated by PCR-based site-directed mutagenesis using oligonucleotides and templates listed in Table S1 and inserted into the SalI site of pHE-C or phsp-GFP-Halo as described above. All of the plasmids were confirmed by DNA sequencing.

### Transfection, infection, and microscopy

Plasmid transfection and virus infection were performed as described previously (4). HaloTag staining was conducted with the HaloTag TMR Ligand according to the manufacturer’s instructions (Promega). Confocal images were acquired with a Zeiss ConfoCor 2 (63×, 1.4 NA, oil) using a 488-nm laser line for GFP and a 543-nm laser line for tetramethylrhodamine (TMR). All experiments and image acquisition were performed on live cells.

### Image analysis

Image processing was performed using Fiji (19) with the extracellular background intensity subtracted from each image. Compartment fluorescence was calculated as the mean intensity of pixels in the nucleus, cytoplasm, virogenic stroma, or nucleoplasm by using manual image segmentation. FRET experiments used GFP and TMR channel images obtained with 488 nm-laser irradiation. FRET images were generated in Fiji based on the image calculation: TMR / (GFP + TMR). FRET indices were calculated from the FRET images as the mean intensity of pixels in the nucleus, cytoplasm, virogenic stroma, or nucleoplasm as determined by manual image segmentation.

### Focus formation assay

The presence of the IE1 focus was manually scored under the microscope with varying focus according to cell thickness. For each experiment, more than 100 randomly selected cells were analyzed and mean indices were calculated from three independent experiments.

## ACKNOWLEDGMENTS

We thank J. Joe Hull for critical reading of the manuscript. This research was supported in part by JSPS KAKENHI Grant Number 17K08162.

## References

1. G. F. Rohrmann, Baculovirus Molecular Biology. National Center for Biotechnology Information (US) 4th edition (2019).

2. T. Nagamine, Apoptotic arms races in insect baculovirus coevolution. Physiol. Entomol. 47, 1–10 (2022).

3. K. Okano, V. S. Mikhailov, S. Maeda, Colocalization of baculovirus IE-1 and two DNA-binding proteins, DBP and LEF-3, to viral replication factories. J. Virol. 73, 110–119 (1999).

4. Y. Kawasaki, S. Matsumoto, T. Nagamine, Analysis of baculovirus IE1 in living cells: dynamics and spatial relationships to viral structural proteins. J. Gen. Virol. 85, 3575–3583 (2004).

5. T. Nagamine, A. Abe, T. Suzuki, N. Dohmae, S. Matsumoto, Co-expression of four baculovirus proteins, IE1, LEF3, P143, and PP31, elicits a cellular chromatin-containing reticulate structure in the nuclei of uninfected cells. Virology 417, 188–195 (2011).

6. V. A. Olson, J. A. Wetter, P. D. Friesen, The highly conserved basic domain I of baculovirus IE1 is required for hr enhancer DNA binding and hr-dependent transactivation. J. Virol. 77, 5668–77 (2003).

7. T. Nagamine, Y. Kawasaki, T. Iizuka, S. Matsumoto, Focal distribution of baculovirus IE1 triggered by its binding to the hr DNA elements. J. Virol. 79, 39–46 (2005).

8. S. Kosugi, M. Hasebe, M. Tomita, H. Yanagawa, Systematic identification of yeast cell cycle-dependent nucleocytoplasmic shuttling proteins by prediction of composite motifs. Proc. Natl. Acad. Sci. USA 106, 10171–10176 (2009).

9. L. He et al., Systematic analysis of nuclear localization of Autographa californica multiple nucleopolyhedrovirus proteins. J. Gen. Virol. 102, 001517 (2021).

10. V. A. Olson, J. A. Wetter, P. D. Friesen, Oligomerization mediated by a helix-loop-helix-like domain of baculovirus IE1 is required for early promoter transactivation. J. Virol. 75, 6042–51 (2001).

11. V. A. Olson, J. A. Wetter, P. D. Friesen, Baculovirus transregulator IE1 requires a dimeric nuclear localization element for nuclear import and promoter activation. J. Virol. 76, 9505–15 (2002).

12. F. Diella et al., Understanding eukaryotic linear motifs and their role in cell signaling and regulation. Front. Biosci. 13, 6580–603 (2008).

13. N. E. Davey, G. Travé, T. J. Gibson, How viruses hijack cell regulation. Trends. Biochem. Sci. 36, 159–69 (2011).

14. G. R. Kovacs, J. Choi, L. A. Guarino, M. D. Summers, Functional dissection of the Autographa californica nuclear polyhedrosis virus immediate-early 1 transcriptional regulatory protein. J. Virol. 66, 7429–7437 (1992).

15. J. M. Slack, G. W. Blissard, Identification of two independent transcriptional activation domains in the Autographa californica multicapsid nuclear polyhedrosis virus IE1 protein. J. Virol. 71, 9579–87 (1997).

16. A. Kondo, S. Maeda, Host range expansion by recombination of the baculoviruses Bombyx mori nuclear polyhedrosis virus and Autographa californica nuclear polyhedrosis virus. J. Virol. 65, 3625–3632 (1991).

17. T. Nagamine, T. Inaba, Y. Sako, A nuclear envelop-associated baculovirus protein promotes intranuclear lipid accumulation during infection. Virology 532, 108–117 (2019).

18. K. A. Motohashi, Simple and efficient seamless DNA cloning method using SLiCE from Escherichia coli laboratory strains and its application to SLiP site-directed mutagenesis. BMC Biotechnol. 15, 47 (2015).

19. J. Schindelin et al., Fiji: an open-source platform for biological-image analysis. Nat. methods 9, 676–682 (2012).

